# ReprDB and panDB: minimalist databases with maximal microbial representation

**DOI:** 10.1101/214916

**Authors:** Wei Zhou, Nicole Gay, Julia Oh

## Abstract

**Background:** Profiling of shotgun metagenomic samples is hindered by a lack of unified microbial reference genome databases that i) assemble genomic information from all open access microbial genomes, ii) have relatively small sizes, and iii) are compatible to various metagenomic read mapping tools. Moreover, computational tools to rapidly compile and update such databases to accommodate the rapid increase in new reference genomes do not exist. As a result, database-guided analyses often fail to profile a substantial fraction of metagenomic shotgun sequencing reads from complex microbiomes.

**Results:** We report pipelines that efficiently traverse all open access microbial genomes and assemble non-redundant genomic information. The pipelines result in two species-resolution microbial reference databases of relatively small sizes: reprDB, which assembles microbial representative or reference genomes, and panDB, for which we developed a novel iterative alignment algorithm to identify and assemble non-redundant genomic regions in multiple sequenced strains. With the databases, we managed to assign taxonomic labels and genome positions to the majority of metagenomic reads from human skin and gut microbiomes, demonstrating a significant improvement over a previous database-guided analysis on the same datasets.

**Conclusions:** reprDB and panDB leverage the rapid increases in the number of open access microbial genomes to more fully profile metagenomic samples. Additionally, the databases exclude redundant sequence information to avoid inflated storage or memory space and indexing or analyses time. Finally, the novel iterative alignment algorithm significantly increases efficiency in pan-genome identification and can be useful in comparative genomic analyses.

## Background

The microbiome field has been revolutionized by sequencing technologies that enable reconstruction of microbial community composition and function. Metagenomic whole-genome shotgun sequencing (mWGS) samples the full genomic complement of a community to provide a high-resolution reconstruction of its species, strains, and even single-nucleotide polymorphisms. With adequate sequencing depth, mWGS data contain information on the scale of billions of short reads that can be deconvoluted to generate compositional and functional profiles of the hundreds to thousands of microbial species existing in a given microbiome.

Extracting information such as species identities from this vast body of intrinsically complex data is of great biological interest, yet methodologically challenging [1]. To accommodate this data type, methods to analyze mWGS data require high sensitivity (the proportion of data that can be interpreted), high specificity (the proportion of data that are correctly interpreted) and high speed. Commonly, mWGS data are assigned a taxonomic or functional label based on their alignment to a most plausible genome position in a reference database containing microbial genome sequences (for example, see [2-4]). Thus, the sensitivity, specificity, and speed of such database-guided analyses all depend on the intrinsic qualities of the reference database.

For optimal utility, a reference database should have maximal comprehensiveness, which maximizes the sensitivity and specificity of sequence mapping, and minimal redundancy, which minimizes storage space, memory space, and analysis time. Moreover, an ideal reference database should have the ability to accommodate the extensive intra-species genetic diversity, or “pan-genome” [5], that is unique to sub-species or strains within a species and is poorly captured by current databases. To the best of our knowledge, no microbial reference databases have been constructed with the considerations of maximal comprehensiveness and minimal redundancy. Consequently, analyses that utilize current reference databases often fail to align a substantial fraction of mWGS reads. For example, more than 40% of human skin mWGS reads and 60% of human stool mWGS reads remained unmapped to any genomic positions when analyzed against a reference database that combined the Human Microbiome Project, and manually selected prokaryotic and fungal genomes from Refseq that are present on human skin [4]. Other studies have confirmed these estimates; for example, 58%±2.2% of human gut species richness was estimated to be uncharacterizable with a different prokaryotic reference database [6]. Therefore, it is likely that many database-guided analyses have significantly underestimated sample biodiversity and, in turn, biased conclusions drawn from comparisons of different experimental cohorts.

Current efforts to improve metagenomic profiling have focused primarily on new indexing and searching algorithms, which could synergistically benefit from reference databases with improved comprehensiveness and minimal redundancy. For example, bioinformatics tools such as Kraken [7] and Livermore Metagenomics Analysis Toolkit (LMAT) [8] use k-mer-based search algorithms to achieve rapid taxonomic classification. The primary advantage of such approaches is that, theoretically, they can flexibly use different reference databases, including all complete and draft microbial genomes in Genbank, yet the usage of such massive databases strongly inflates storage or memory space and indexing or searching time. Moreover, other analytical frameworks that are incompatible with these searching algorithms (e.g., probabilistic read assignment enabled by Pathoscope [9, 10]) could also benefit significantly from improved databases. On the other hand, tools such as MetaPhlan [11, 12] that do emphasize database quality limit their usage to very specific tasks (such as compositional estimation of a whole sample) by only searching taxonomically informative genome regions and leaving the majority of the metagenomic reads unclassified.

In general, rapid sequence classification algorithms trade memory and storage space for speed by building a substantially large index of the database that is easy to search against. However, this strategy prohibits such methods from incorporating new genomics data, which is problematic given the exponential increase in the number of genomes sequenced in recent years. For instance, LMAT created a 500 GB k-mer index of less than 5,000 microbial species in 2011 [8]; however, at the time of the drafting of this manuscript, GenBank had accumulated genome assemblies of over 80,000 microbial strains from over 20,000 species. The dramatic growth in draft microbial genomes challenges our ability to compile and update reference genome databases in a manner that maintains their compatibility to various bioinformatics tools and utility to the metagenomics community.

To address these limitations, we sought to assemble high-quality genome databases that are compatible with various indexing, searching and analytical algorithms. Here, we describe two new pipelines that each efficiently compiles distinct non-redundant, species-resolution reference databases comprising all open access microbial genomes: database “reprDB” is compiled from microbial representative or reference genome sequences, and database “panDB” consists of the pan-genome sequences of known microbial species, for which we developed an iterative alignment algorithm that efficiently extracts the pan-genomic sequences from a set of conspecific strain genomes. We demonstrate that these databases have the following advantages: they 1) can be re-compiled automatically and efficiently, 2) provide species-level resolution while including strain information, 3) are limited in size, and 4) are able to assign taxonomic labels to the majority of genomic sequence data from human microbiomes.

## Results

### Properties of reprDB and panDB

To more effectively leverage the rich knowledge base of sequenced genomes without excessive analyses time and space requirements, scalable algorithms are required to compress the redundant sequence information in a given database. In particular, conspecific microbial strains contain very similar genomic regions that can be compressed into one representative sequence to greatly reduce the size of the database. For reprDB, intraspecific redundancy is removed by including the representative and reference genomes for each microbial species while discarding the rest of the strain genomes of that species. Therefore, reprDB retains species-level resolution and has minimal size, including 7,018 bacterial, 339 archaeal, 790 fungal and 7,035 viral species totaling 57GB in plain text format.

While reprDB has minimal size and retains species-level resolution, it does not capture intraspecific diversity, which often contains a significant fraction of genetic information in microbial communities. To complement reprDB, we developed panDB with the goal of incorporating sequence information from all conspecific strains, but in a non-redundant fashion. However, identification of non-redundant genomic regions among conspecific strains conventionally requires multiple whole-genome alignment (Figure 1A) that relies on polynomial-time algorithms to compare all aligned genomes and identify collinear genomic regions, which is excessively time-consuming to cover all available microbial species. Therefore, we designed iterative alignment (Figure 1B), a new and scalable algorithm suitable to extract non-redundant pan-genome information from a set of closely related genome sequences. Iterative alignment efficiently assembled panDB comprising pan-genome sequences of 13,485 bacteria, 676 archaea, 864 fungi and 5,578 viruses with a total size of 87GB in plain text format.

**Figure 1.**
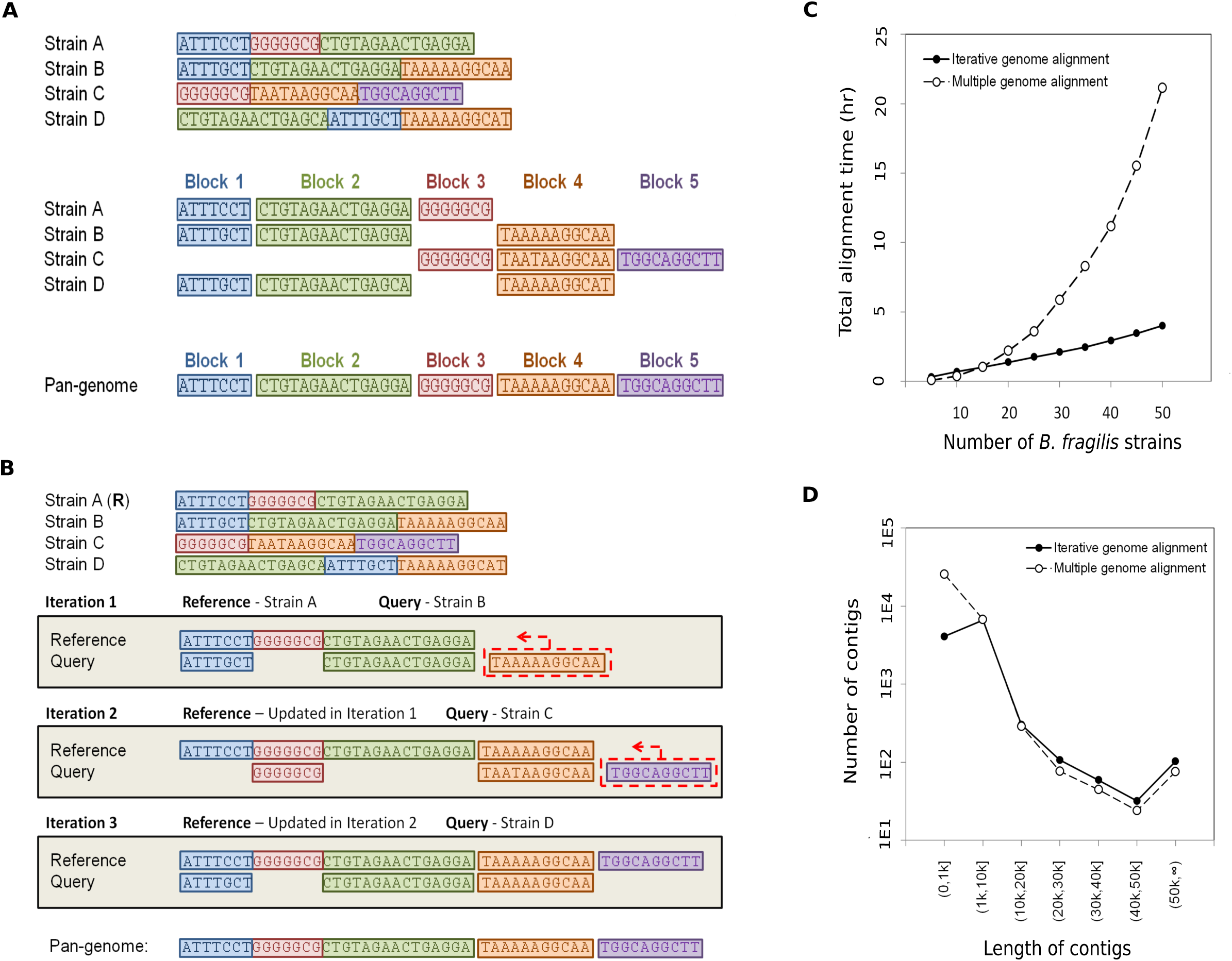
The iterative alignment algorithm. 1A, conventional multiple whole-genome alignment identifies similar genomic regions (blocks) among genome sequences; the union of all block sequences represents the species pan-genome. 1B, the iterative alignment algorithm is demonstrated with a microbial species having four sequenced strains (strain A-D) where strain A is selected as the representative genome, denoted by (**R**). In the first alignment iteration, a query genome is selected (strain B) and aligned to the reference (strain A), identifying the genomic region that is present in the query but not in the reference (orange region). The orange region is then appended to the reference genome to generate an updated version. In the next iteration, a new query genome (strain C) is aligned with the updated reference genome. The query-exclusive purple region is identified and appended to the reference genome to generate a new updated version. The iteration continues until all strains are considered. The final reference represents the species pan-genome. 1C, the total alignment time for processing different numbers of *B. fragilis* strains using multiple whole-genome alignment or iterative alignment. 1D, the distribution of contig sizes that constitute the *B. fragilis* pan-genome identified using multiple whole-genome alignment or iterative alignment.

### Iterative alignment

Our iterative alignment algorithm traverses a list of strain genomes to identify and concatenate non-redundant genomic regions (Figure 1B). It purposefully avoids the computationally-expensive multiple whole-genome alignments, and extracts more contiguous pan-genome sequences than whole-genome alignments. Iterative alignment dramatically increased the efficiency of pan-genome sequence extraction compared to multiple whole-genome alignment (Table 1). Iterative alignment has an empirical time complexity that scales approximately linearly with the number of genomes with a theoretical worst-case complexity of *O(n*^*2*^) when there is completely no alignable region between any pair of genomes, which is significantly more efficient than the polynomial-time multiple whole-genome alignment when the number of genomes is large (Figure 1C). In addition to the remarkable increase in speed, iterative alignment only adds contigs of genomic regions to the growing reference sequence, but does not segment the contigs that are already added to the reference sequence. This property of iterative alignment results in significantly longer contigs for a less fragmented pan-genome per species (3,524 bp versus 1,238 bp per contig on average for *B. fragilis*) (Figure 1D) than conventional multiple whole-genome alignment, while retaining very similar pan-genome sizes (39.9 Mbp versus 39.5 Mbp for *B. fragilis*).

**Table 1.** Alignment time of strain genomes using MWGA and Iterative alignment.

| Species | Number of aligned strains | Alignment time |  |
| --- | --- | --- | --- |
|  |  | MWGA<br>(hours) | Iterative alignment<br>(hours) |
| *Staphylococcus epidermidis* | 50 | 4.5 | 3 |
| *Enterobacter cloacae* | 50 | 53 | 7 |
| *Agrobacterium tumefaciens* | 25 | 19.5 | 2 |
| *Burkholderia mallei* | 50 | 38 | 5.5 |
| *Gilliamella apicola* | 48 | 3 | 2 |
| *Bacteroides fragilis* | 50 | 21.5 | 4 |
| *Mycobacterium abscessus* | 50 | 15.5 | 4 |
| *Clostridium botulinum* | 50 | 28 | 4 |

### Identification of assemblies with exogenous genomic sequences

Microbial genome assemblies in Genbank could contain exogenous sequences, either due to lateral transfer or sample contamination. To account for this possibility, we have included a pipeline in our GitHub repository that detects potentially exogenous contigs in the pan-genome sequences in panDB. To demonstrate the usage of the pipeline, we searched for pan-genomes in panDB that contain *Escherichia coli*-like sequences by aligning a set of 81 *E. coli*-specific marker genes extracted from the MetaPhlAn database to panDB [11, 12]. Cut-offs of alignment summary statistics, such as sequence identifies, e-values and coverage of the aligned region are adjustable by the user. For demonstration, a total of 273 species pan-genomes align to at least one *E. coli*-specific marker gene with sequence identities over 0.9 and e-values less than 10^-6^. Most of these pan-genomes contain only one fragmental alignment to the marker genes (*i.e.* the aligned region covers less than 50% of the marker gene sequence), which could have resulted from true homology of gene motifs instead of exogeneous *E. coli* sequences. 68 of the 273 pan-genomes have contigs that either align to at least two marker genes, or align to at least one marker gene while the aligned region covers more than 50% of the marker gene sequence. Among the 68 pan-genomes, the majority (73%) are from the genera *Escherichia* and *Shigella* (Figure 2), which are phylogenetically close to *E. coli*. The rest of the species pan-genomes are evolutionarily distant from *E. coli* yet still contain genomic regions highly similar to the *E. coli* marker genes. These pan-genomes likely contain contaminated or laterally transferred *E. coli* sequences, while they constitute less than 0.1% of all pan-genomes in panDB.

**Figure 2.**
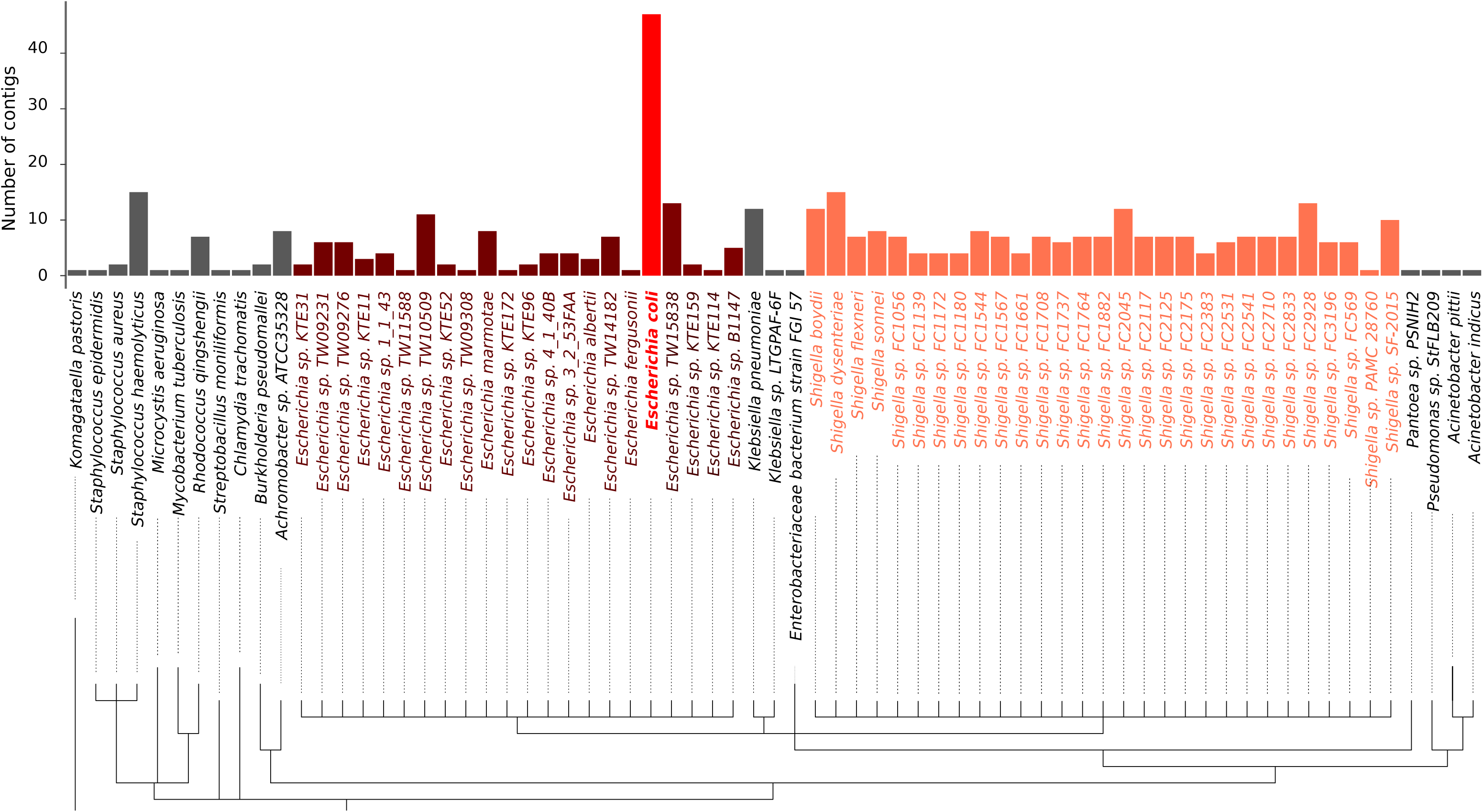
Species pan-genomes containing regions highly similar to *E. coli*. The number of contigs that either align to at least two marker genes, or align to at least one marker gene while the aligned region covers more than 50% of the marker gene sequence are shown for each species pan-genome.

### Read classification of in silico synthetic communities

To test the ability of reprDB and panDB to classify metagenomic reads at different taxonomic resolutions, from communities of common and uncommon species, and from communities of different levels of complexity, we classified sequencing reads sampled from three types of synthetic communities using a short read aligner (Bowtie 2 [13]) and a read classifier (Pathoscope 2.0 [9]).

We first classified sequencing reads simulated from three classes of low-complexity *in silico* synthetic communities: 1) communities consisting of 5, 10, or 20 strains from the same species (either *Staphylococcus epidermidis* or *Bacteroides fragilis*), 2) communities consisting of 5 species from the same genus (either *Staphylococcus* or *Bacteroides*), and 3) communities consisting of 5 species representing 5 different phyla. Because our databases treat species as the lowest taxonomic unit, the synthetic communities composed of conspecific strains were only used to test mapping sensitivity. ReprDB and panDB were able to classify the majority (over 80% for reprDB and over 98% for panDB) of reads from the communities consisting of different numbers of *S. epidermidis* or *B. fragilis* strains (Figure 3A). The phylum-level read classification is highly accurate: over 97% and 99.9% of all reads were classified to the correct phylum by reprDB and panDB respectively (Figure 3B). Species-level read classification is also considerably accurate: reprDB correctly classified 86% and 98%, while panDB correctly classified 92% and 77% of the simulated reads from *Bacteroides* and *Staphylococcus* species, respectively (Figure 3B). The majority of the incorrectly assigned *Staphylococcus* reads were mapped to *Staphylococcus haemolyticus* (20.5% of total reads were assigned to *S. haemolyticus* after Pathoscope reassignment and 13.9% before Pathoscope reassignment). As *S. haemolyticus* is not present in the synthetic community but has a very large pan-genome (22Mbp), this skews the estimated parameter values in the Bayesian read assignment model of Pathoscope 2.0. The sensitivity and specificity of read assignment is robust to the number of simulated reads (Figure 3B) which represents the difference in sequencing depth (1.6-3.9x coverage for 500,000 reads and approximately 3.3-7.7x coverage for 1,000,000 reads).

**Figure 3.**
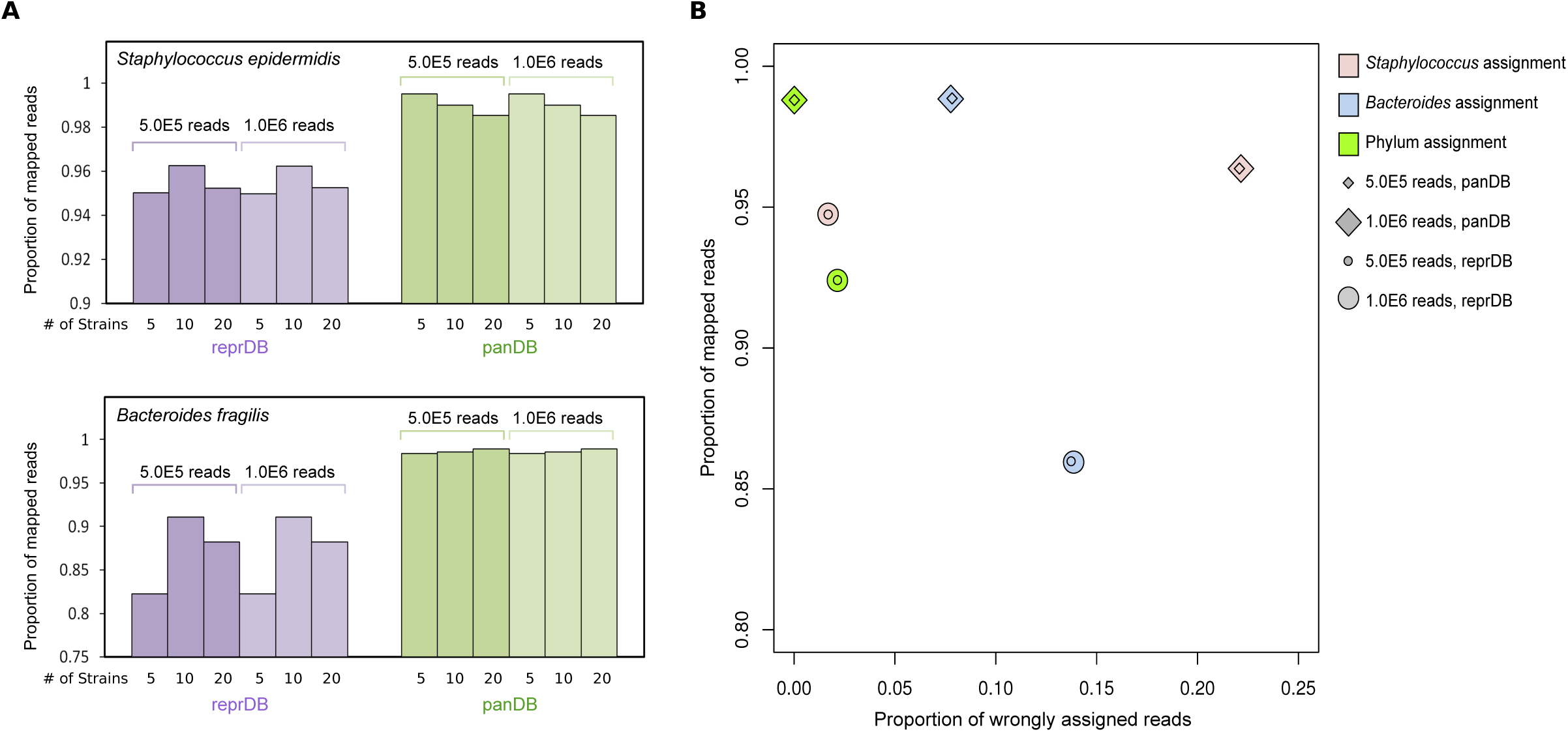
Read classification of low-complexity *in silico* synthetic communities using Bowtie 2 and Pathoscope 2.0. 3A, the proportion of mappable reads generated from synthetic communities composed of 5, 10 or 20 *S. epidermidis* or *B. fragilis* strains. 3B, the proportion of mappable versus wrongly assigned reads generated from synthetic communities composed of 5 different *Staphylococcus* or *Bacteroides* species, or species representing 5 different bacterial phyla.

Next, to test the ability of reprDB and panDB to classify reads from a low-complexity community composed of common bacterial species, we analyzed a mock metagenome community composed of 21 evenly mixed bacterial strains from ‘mockrobiota’, originally contributed by Kozich et al [14, 15]. Both reprDB and panDB can classify 96.0% of the total reads (Figure 4A and 4B). Based on reprDB, 92.9% of the reads were mapped to one of the 21 species that constitute the community (Figure 4A), and 93.1% of the reads were mapped to one of the 18 genera in the community (Figure 4B). As anticipated, the 21 species have approximately even abundances based on reprDB (Pielou’s evenness index = 0.83). However, for panDB, a significantly fewer proportion of reads (53.5%) were correctly mapped to the 21 species, and the estimated relative abundances are less even (Pielou’s evenness index = 0.525) (Figure 4A). Based on the fact that panDB correctly assigned a relatively high proportion of reads (87.5%) to the 18 genera that constitute the community (Figure 4B), it can be deduced that panDB mistakenly assigned a subset of reads to a closely related species pan-genome in the same genus, similar to what was observed when assigning the *Staphylococcus* reads in the previous analyses. The results suggest that reprDB can accurately profile communities composed of common bacteria, while panDB is less accurate at species-level profiling due to the interference of closely related species pan-genomes.

**Figure 4.**
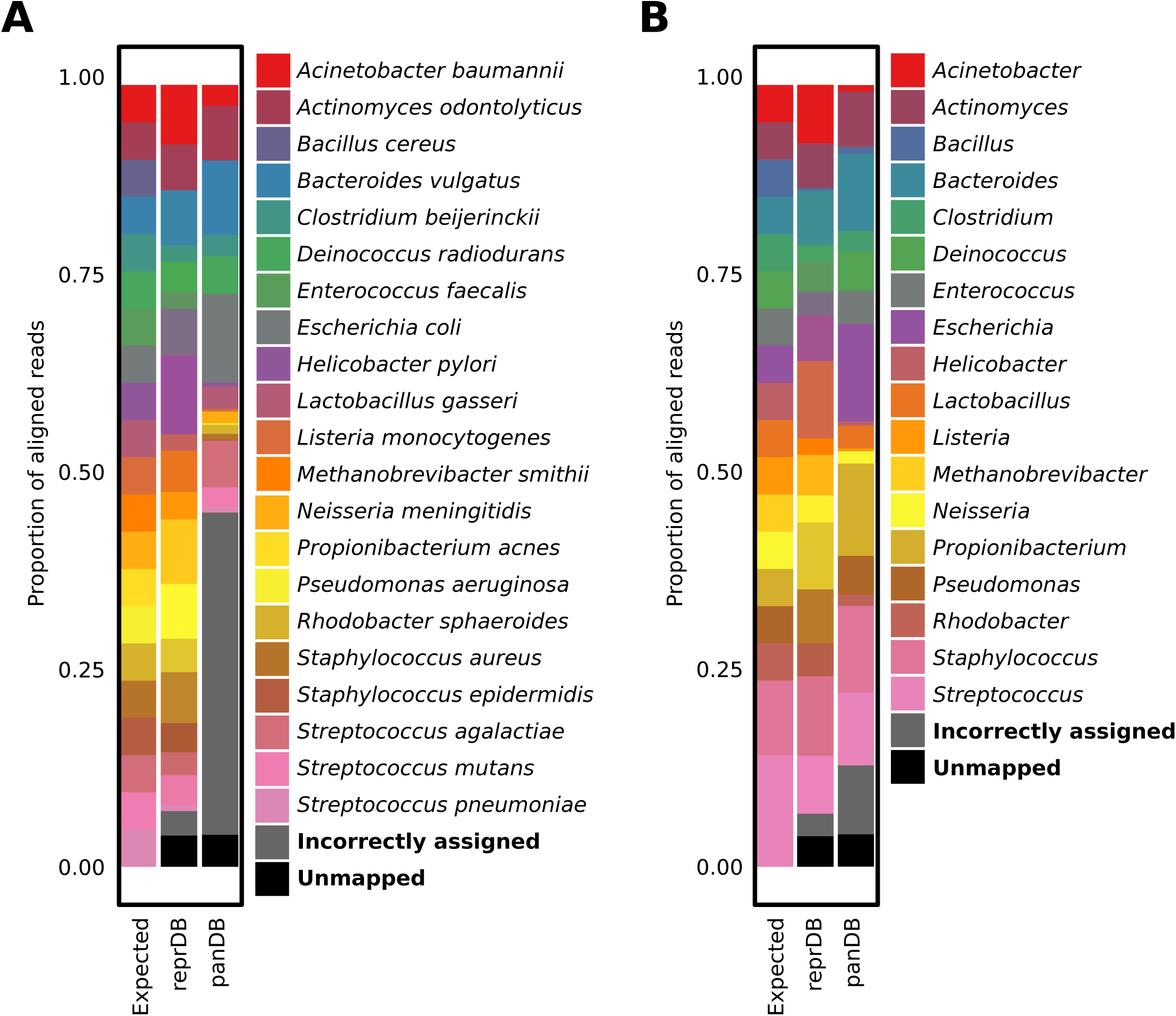
Read classification of the mockrobiota mock community using Bowtie 2 and Pathoscope 2.0. 4A, the proportion of reads mapped to the microbial species present in the mock community. Bowtie 2 was used for read alignment and reads that aligned to multiple genome locations were assigned to the most probable genome location using Pathoscope 2.0. 4B, the proportion of reads mapped to the microbial genera present in the mock community. Bowtie 2 was used for read alignment and reads that aligned to multiple genome locations were assigned to the most probable genome location using Pathoscope 2.0.

Finally, to demonstrate that including more species pan-genome information in the database could improve profiling of unknown and high-complexity communities, we tested the performance reprDB and panDB using 5 datasets from Critical Assessment of Metagenome Interpretation (CAMI) [16], which contain over 700 predominantly unpublished isolate genomes, including multiple strains per species, microdiversity and non-chromosomal elements [16]. Because most of the genomic sequences lack taxonomic typing, we compared our results to the “gold standard profiling”, which was generated based on the Refseq and NCBI bacterial genome database and can be downloaded from CAMI. ReprDB and panDB were able to characterize an average of 49.7% and 75.1% of all reads in the datasets – approximately 74% and 163% more reads compared to the gold standard profiling (Figure 5A and 5C). Moreover, although reprDB and panDB characterized a much larger subset of the data, the estimated abundances of the taxa still significantly and positively correlate to the gold standard (Figure 5B and 5C, average Pearson correlation coefficient=0.62 between the gold standard and reprDB estimates, and average Pearson correlation coefficient=0.70 between the gold standard and panDB estimates). In addition, the Refseq database used to generate the gold standard profiling contains not only bacteria genomes, but also plasmid and gene sequences, and is much larger in size (37.6GB of bacterial genome sequences and 482.8GB of non-genome sequences) than reprDB and panDB, but it only classified a much smaller subset of reads in the datasets compared to reprDB and panDB (Figure 5A and 5C). The results suggest that reprDB and panDB are informative and compact, while panDB is especially powerful when characterizing highly complex communities with strain-level diversity and potentially unknown microbial species.

**Figure 5.**
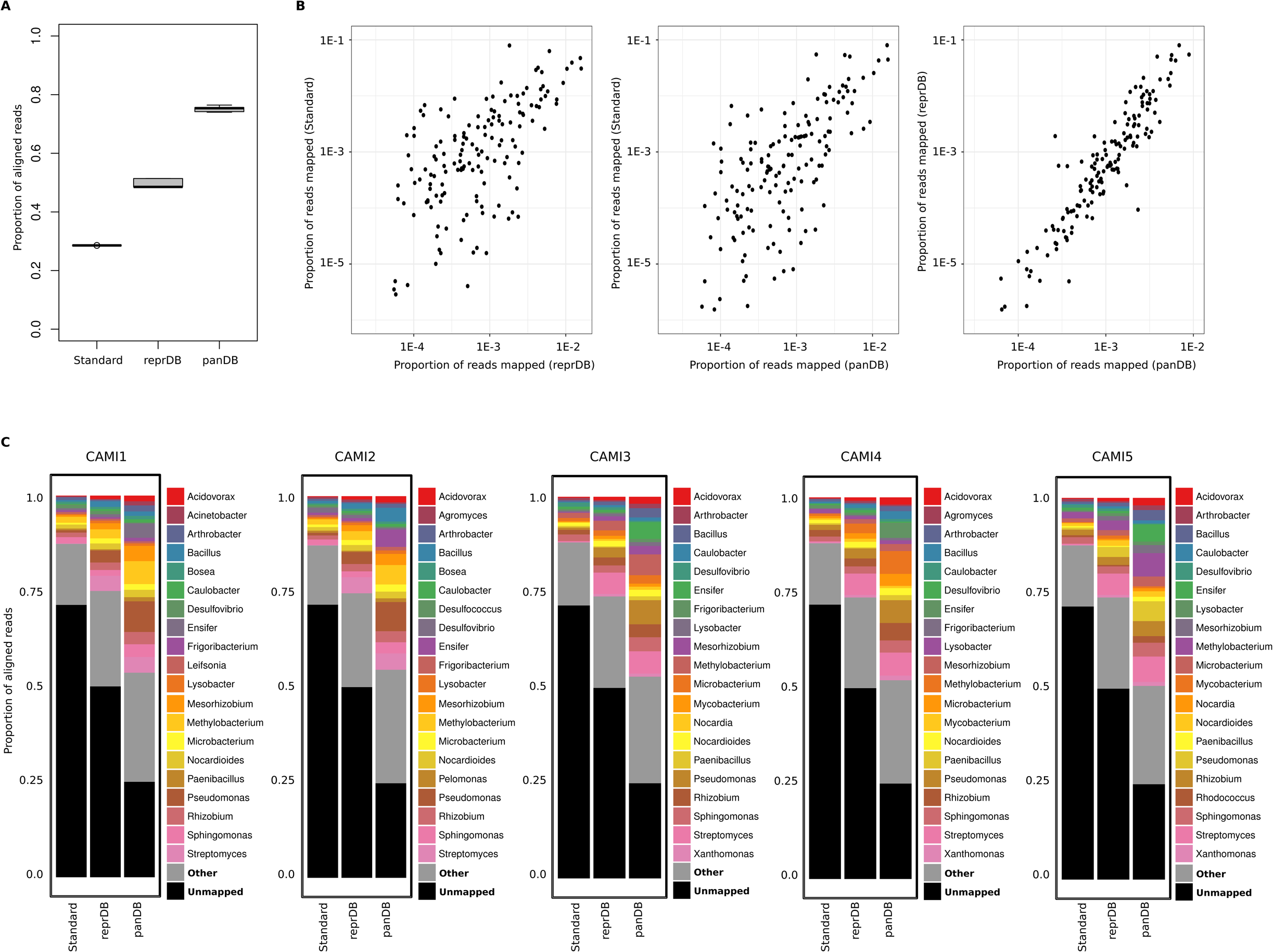
Read classification of high-complexity synthetic communities from CAMI using Bowtie 2 and Pathoscope 2.0. 5A, the proportion of reads that were classified using reprDB, panDB and the gold standard profiling provided by CAMI (labeled as “Standard”). 5B, the proportions of reads from the mapped to different microbial genera based on reprDB, panDB and the gold standard profiling provided by CAMI (labeled as “Standard”). 4C the proportion of reads mapped to the 20 most abundant microbial genera characterized based on reprDB, panDB and the gold standard profiling provided by CAMI (labeled as “Standard”).

### Compatibility with Kraken

To show that reprDB and panDB is compatible with read classification tools other than Pathoscope 2.0, we used Kraken to classify sequencing reads based on reprDB, panDB and the standard Kraken library [7]. In contrast to Pathoscope 2.0, which assigns each read to a single genome, Kraken assigns each read to its lowest taxonomic level [7]. Using the *in silico* communities synthesized in this study, we assessed the total proportion of reads classified based on the databases. A similar proportion of reads sampled from the *S. epidermidis* strains were classified based on reprDB, panDB, and the standard Kraken database, while panDB classified more reads sampled from *B. fragilis* strains than reprDB or the standard Kraken database (Figure 6A). The results were consistent to the observation when reads were classified using Pathoscope 2.0 (Figure 3A). Next, we assessed the consistency between the genome from which a read was sampled and the taxonomic node to which the read was assigned to. panDB exhibits the highest consistency in read classification (Figure 6B). 25% of the reads generated from the community composed of 5 *Bacteroides* species was not correctly assigned using reprDB, and an even larger amount (34%) failed to be correctly assigned using the standard Kraken database (Figure 6B). These observations were consistent to the results generated using Pathoscope 2.0 (Figure 3B). While Pathoscope 2.0 incorrectly assigned a significant proportion of multi-mapped *Staphylococcus* reads to *S. haemolyticus* based on panDB (Figure 3B), Kraken did not exhibit this problem for it classifies multi-mapped reads to the lowest common ancestral node [7].

**Figure 6.**
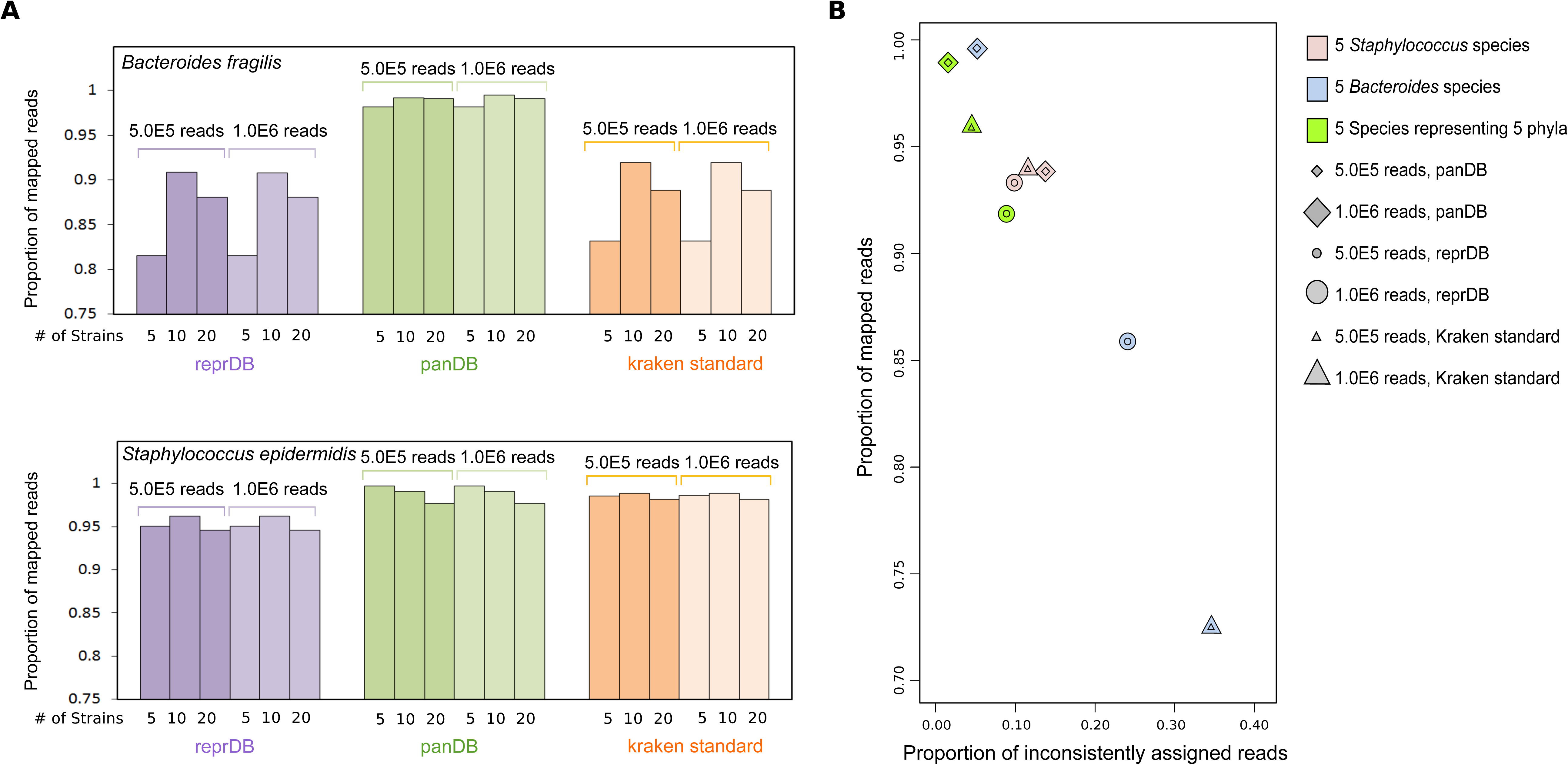
Read classification of low-complexity *in silico* synthetic communities using Kraken. 6A, the proportion of mappable reads generated from synthetic communities composed of 5, 10 or 20 *S. epidermidis* or *B. fragilis* strains. 6B, the proportion of mapped reads versus inconsistently classified reads generated from synthetic communities composed of 5 different *Staphylococcus* or *Bacteroides* species, or species representing 5 different bacterial phyla. “Inconsistent” means the read was neither assigned to the species from which it was sampled, nor assigned to the an ancestral taxon of the species from which it was sampled.

### Read classification of the skin and stool mWGS samples

We then re-analyzed the mWGS data from over 800 human skin and stool microbiome samples described in Oh et al. [4] to determine if these databases could provide new insights into the species composition of metagenomic samples. ReprDB characterized 66.0±18% of the skin mWGS reads (Figure 7A, left panel) and 73.3±9% of stool mWGS reads (Figure 7A, right panel), in comparison to 58.8±24% and 33.8±14%, respectively, in the initial publication [4]. PanDB, with its inclusion of as many non-redundant genome regions as possible, classified an even larger fraction of reads (71.6±18% and 80±10%, respectively) (Figure 7A). Consistent with previous reports [4], *Propionibacterium, Corynebacterium* and *Staphylococcus* were observed as the most abundant bacteria genera in the skin samples (Figure 7B, upper panel), while *Bacteroides* was observed as the most abundant in the stool samples (Figure 7B, lower panel). A substantial amount of reads mapped to uncultured species—species that have no axenic culture for formal description—based on the panDB, especially in the stool samples (Figure 7B).

**Figure 7.**
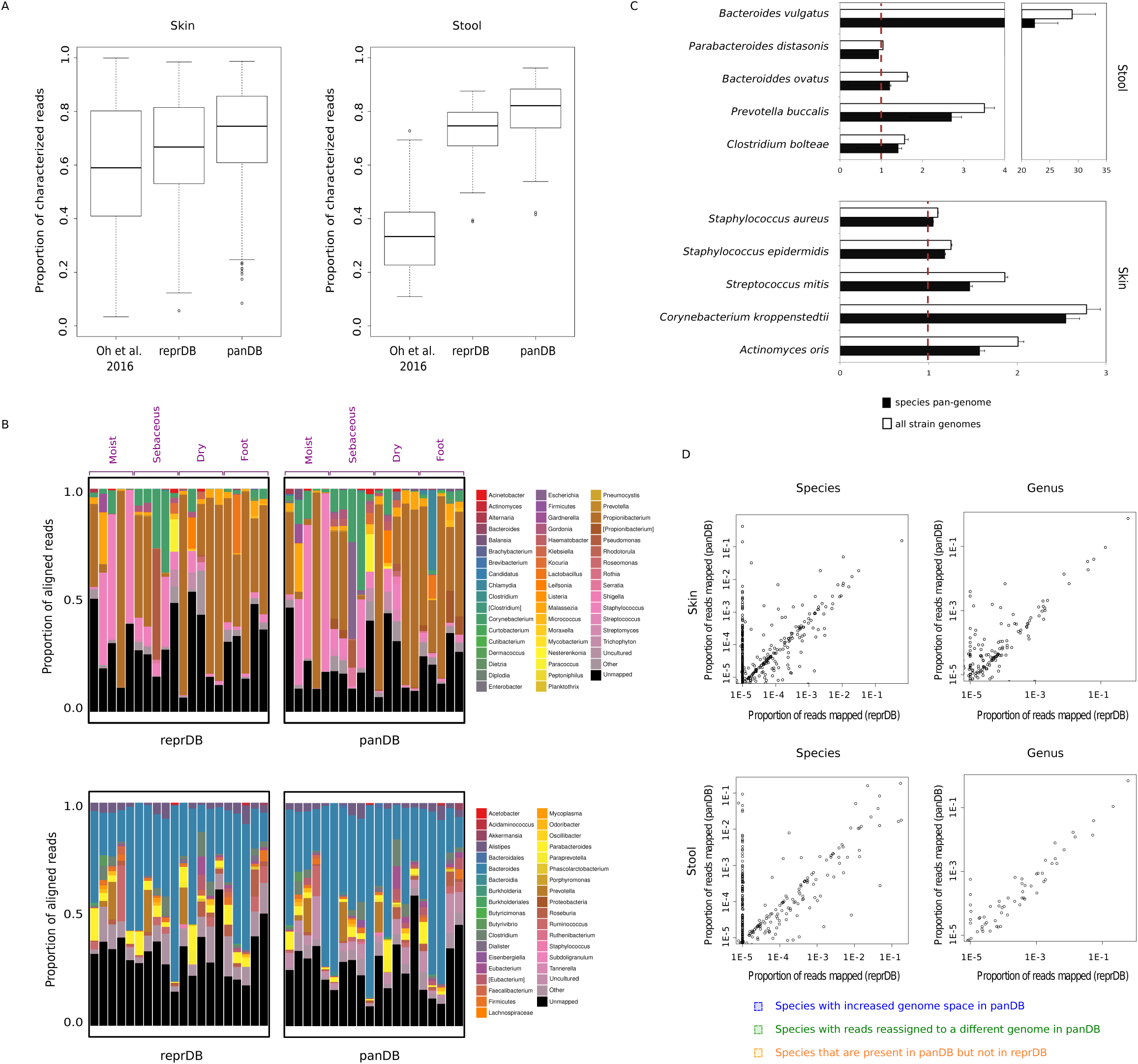
Read classification of skin and stool mWGS samples. 7A, the proportion of skin (left) and stool (right) reads that are classified using reprDB, panDB and a previous database described in [4]. 7B, the most abundant microbial groups in selected skin (upper) and stool (lower) samples based on reprDB and panDB. Microbial species are grouped by their lowest taxonomic level at or above genus level. Tentative classifications and misplaced classifications that are waiting for revisions are shown in square brackets according to the convention of the NCBI Taxonomy database [39]. Uncultured species are classified into an independent group [40]. 7C, the proportion of reads mapped to the pan-genome or all sequenced strain genomes of selected stool (upper) and skin (lower) species, normalized to the proportion of reads mapped to the representative genome(s) of the same species (dashed line). 7D, the proportions of reads aligned to different genera (right) or species (left) based on reprDB (x axis) and panDB (y axis). Results are shown for a randomly selected skin sample (MET0284, upper) and a random stool sample (SRS011061, lower). In the upper left panel, three different types of species that recruit inconsistent proportion of reads based on reprDB and panDB are shown in colored boxes. Blue box: species with increased genome space in panDB; green box: species with reads reassigned to a different genome in panDB; orange box: species that are present in panDB but not in reprDB.

To assess how well these databases capture the within-species diversity observed in the skin and stool samples, we compared the fraction of reads that aligned to the representative genome, the pan-genome sequence, and all strain genomes of a given species that are aligned to derive the pan-genome sequence. For this purpose, we randomly chose 5 microbial species that have multiple sequenced strain genomes and are relatively abundant in skin or stool samples, respectively (Table 2). Many species contain significant within-species diversity that was not characterized by the representative sequences; in an extreme case, the representative genomes of *B. vulgatus* captured less than 5% of all reads mappable to the species pan-genome (Figure 7C, upper panel). Overall, the pan-genome sequences alone were able to profile a similar fraction of reads with the unaligned strain genomes combined, while having a significantly reduced size (Table 2).

**Table 2.** Sizes of reprDB species representative genomes, panDB species pan-genomes and the total size of sequenced strain genomes used to generate the species pan-genomes.

| Species | Size in reprDB<br>(kbp) | Size in panDB<br>(kbp) | Total size of strain genomes<br>(kbp) |
| --- | --- | --- | --- |
| *Bacteroides vulgatus* | 5,163 | 9,954 | 41,523 |
| *Parabacteroides distasonis* | 4,811 | 9,486 | 53,014 |
| *Bacteroides ovatus* | 6,791 | 14,112 | 69,311 |
| *Prevotella buccalis* | 6,010 | 3,886 | 8,854 |
| *Clostridium bolteae* | 5,432 | 9,349 | 45,990 |
| *Staphylococcus aureus* | 14,177 | 12,080 | 239,676 |
| *Staphylococcus epidermidis* | 5,209 | 17,271 | 153,889 |
| *Streptococcus mitis* | 2,147 | 11,132 | 112,321 |
| *Corynebacterium kroppenstedtii* | 2,447 | 2,715 | 10,197 |
| *Actinomyces oris* | 2,949 | 3,421 | 8,957 |

### Consistency in read assignments based on the two databases

To compare the performances of each database, we compared the taxonomic classification of skin and stool mWGS reads using reprDB and panDB. The fractions of reads assigned to the same microbial genus were generally comparable based on reprDB and panDB, with an average Pearson correlation coefficient of 0.943 for skin samples and 0.958 for stool samples (Figure 7D). However, inconsistency in read assignment increased at the species level, resulting in an average correlation coefficient of 0.323 for skin samples and 0.720 for stool samples (Figure 7D, left panel). Many reads were mapped to species that are present in panDB, but are not included in reprDB because the species have no designated representative genomes (for example, the species data points in the orange square in the upper left panel of Figure 7D). In addition, panDB was able to recruit additional reads to the pan-genome regions of many species compared to reprDB (for example, the species data points in the blue square in the upper left panel of Figure 7D). Moreover, reads that initially mapped to a reprDB species may align to a different genomic region in panDB, consequently decreasing the observed abundance of the species which the reads originally mapped to (for example, the species data points in the green square in the upper left panel of Figure 7D). While reprDB is smaller and more balanced in that it does not over-represent species that have more sequenced strain genomes, panDB provides a more comprehensive classification that incorporates intraspecific diversity.

### Consistency in taxonomic profiles generated based on 16s rRNA and mWGS data

Finally, we compared the taxonomic profiles generated based on reprDB and panDB to the genus-level taxonomic profile generated based on 16s rRNA sequencing for 57 stool samples that have both mWGS and 16s rRNA sequencing data [17]. 16s rRNA sequencing identified 118±47 microbial genera, out of which 45±7% were not identified using either reprDB or panDB based on the paired mWGS data, partly because no whole genome sequences are available in Genbank or Refseq for some of these genera. On the other hand, 91±5% of the 873±341 genera identified using either reprDB or panDB were not identified by 16s rRNA sequencing. The 64±22 genera that are robustly identified by both sequencing methods have highly consistent relative abundance estimates, with an average Pearson correlation coefficient of 0.89 between 16s rRNA and reprDB and 0.88 between 16s RNA and panDB.

## Discussion

Ideally, reference databases for characterizing metagenomic data should be comprehensive but non-redundant to accurately classify as many sequencing reads as possible with maximum efficiency and minimum computational cost. In this study, we described methods to efficiently and automatically compile two microbial genome databases, reprDB and panDB, both of which have manageable sizes and are suitable for analyzing mWGS data. The reference databases we compiled significantly promoted microbial species identification in human-associated microbial communities. For many of the stool and skin microbiome samples tested in this study, close to 100% of the reads were classified using either of the two databases and with high consistency in taxonomic classification, representing a significant improvement over previous analyses [4].

For some samples, however, only a tiny fraction of reads were classified, despite the comprehensive nature of reprDB and panDB. This suggests the presence of microbial clades that have no available genome sequences to date, perhaps owing to generally low prevalence or conditional abundance. Similarly, some of the clades identified by 16s rRNA sequencing were not identified in the shotgun metagenome dataset because no genome sequences were available for these clades, underscoring the need for continued microbial genome discovery and database integration. Alternatively, their lack of representation could result from systematic bias in microbial isolation or sequencing methods. For example, bacteria from the *Prevotella* genus are harder to culture than *Bacteroides* bacteria, which results in significantly more sequenced *Bacteroides* genomes than *Prevotella.* Consequently, significantly fewer *Prevotella* reads can be classified using standard reference databases. A similar bias was also exposed in our simulation study, where a significant number of reads were incorrectly assigned to highly represented and closely related species in the database (Figure 3B and 4A). In general, public sequence databases such as GenBank contain more data for microbial species of clinical, biological, or technical importance, as well as species that are easier to isolate and culture. It therefore follows that databases curated from these sources will propogate this skewing and potentially bias the alignment-based characterization of microbial communities. Therefore, addition of genome sequences of underrepresented yet ecologically important species will make the reference databases not only more informative, but also more balanced and robust for read classification purposes. Given that isolation and sequencing of such species could be technically challenging, this underscores the value of phylogeny-driven sequencing efforts such as GEBA [18-20] as well as innovations in culturomics [21, 22] and single-cell sequencing [23] to improve species representation in reference databases.

ReprDB selectively uses representative or reference genome sequences to minimize its size while providing species-level resolution. Representative or reference genomes are selected from the NCBI RefSeq database based on community consensus, assembly and annotation quality as well as consideration of species-level taxonomic classification, and then manually curated for their metadata by PATRIC. In addition to being compact and high-quality, sequence information in reprDB is more balanced across microbial species, in the sense that the database only includes the representative or reference genomes of a given species, regardless of how many strains have been sequenced for the species. Under circumstances where the closely related species in a community are sparse or skewed, repDB could serve as an important complement to panDB, or even outperform panDB in terms of read classification accuracy especially for commonly observed bacterial species (Figure 3B, 4A and 4B).

On the other hand, panDB purposefully includes as much non-redundant information as possible for each known microbial species in order to classify as many microbial reads as possible. It is moderately larger in size than reprDB and correspondingly classifies a larger fraction of mWGS data. However, this increase is biologically significant because many species possess considerable strain-level diversity within their human-associated habitats, which can only be characterized by accounting for intraspecific diversity in the reference database (Figure 7C and 7D). Moreover, intraspecific genetic variation can differ markedly between species (e.g., 20% of gene families are variable in commensal S. *epidermidis* [24], while 70% are variable in *Salmonella enterica* [25]), requiring an approach that can accommodate large numbers of strains. We show that panDB is especially valuable in characterizing communities of high complexity, with strain-level diversity and with previously unknown species (Figure 5A and 5C), because panDB more comprehensively explores the sequence space that are accessible to each microbial species by accounting for all sequenced strains of that species. Additionally, reads that map to one genome location in reprDB may find a different alignment in panDB. Such relocated reads are not rare, and they can substantially influence read assignment among species. This is because read assignment is commonly inferred under a Bayesian framework in software such as Pathoscope [9] where the assignment of one read influences the probability of the assignment of other reads. It is important to note that we found that read assignments are more precise with panDB, which contains more information than reprDB, but not definitively more accurate due to the above-mentioned bias in representation among species (Figure 3B and Figure 4A). Accuracy of panDB-based read assignment could be further improved by balancing species representation, or introducing read assignment models that explicitly correct for the variation in species representation. Another factor that could influence the accuracy of panDB is the presence of exogenous sequences in genome assemblies, due to either contamination or lateral transfer. Although we show that the proportion of contaminated assemblies is likely small (Figure 2), assemblies containing laterally transferred element are hard to identify computationally based on only the genome sequence. However, when analyzing mWGS datasets, it is possible to minimize the influence of laterally transferred element by combining database-based profiling with *de novo* approaches such as coverage-based binning (for example, see [26]).

The importance of species pan-genomes in analyzing compositional or functional aspects of metagenomic datasets has received much attention. Most methods, however, characterize pan-genomes on the resolution of gene families, ignoring non-coding regions or unannotated coding sequences [27, 28]. This limitation likely arises from both the high computational cost of multiple whole-genome alignment and the lack of alternative algorithms that can efficiently identify pan-genomes based on full genome sequences. Thus, we based panDB on an iterative alignment algorithm which allows rapid extraction of the pan-genome sequence from a set of conspecific strain genomes independent of annotated coding sequences. Iterative alignment is a greedy algorithm, as empirical computation time scales approximately linearly with the number of strain genomes that are aligned (Figure 1C). In addition to superior speed, iterative alignment has two advantages for database compilation. First, the algorithm results in less segmented pan-genome sequences than conventional multiple whole-genome alignment (Figure 1D). During iterative alignment, the reference sequence will only be extended but never trimmed or rearranged. Therefore, the order of bases observed in the reference sequence at any given time, including those bases in the blocks that are appended to the reference sequence, will be preserved throughout the database compilation (Figure 1B). A less segmented pan-genome sequence can reduce the loss of mappable reads that align to the breakpoints between contigs. Second, databases constructed using iterative alignment can be easily updated: each newly added genome can be incorporated into the present pan-genome sequence by conducting one alignment using the pan-genome sequence as the reference and the newly added genome as the query. Thus, panDB is can be repeatedly expanded without fully recompiling the database.

Finally, reference databases that provide high-quality *a priori* knowledge can aid other analytical methods. For example, reprDB and panDB can disentangle mWGS data by partitioning reads according to their species of origin, facilitating fast and accurate assembly of species’ genomes. Comprehensive databases can also improve analysis of genetic polymorphisms in microbial populations by providing more precise alignments for variant base calling. In addition, these databases can be extended to include contextual or sequence-associated information, such as functional annotation of the genome sequences, to reveal new insights of microbial communities with functional relevance. Apart from alignment-based analyses, reprDB and panDB can serve as the benchmark or training set for the construction of predictive models based on sequence features.

## Conclusion

Current microbial reference databases often fail to characterize a large proportion of shotgun metagenomic data from complex microbiomes. We developed efficient algorithms to compile two species-resolution reference databases, ReprDB and panDB, that significantly outperform current databases. ReprDB has minimal size and balances species representation by including reference or representative microbial genomes, while panDB uses a novel algorithm to identify and assemble as much non-redundant sequence information as possible to more fully capture intraspecific genetic diversity. Both databases demonstrate high sensitivity in classifying sequence reads. ReprDB classifies reads from common microbial species with high accuracy, while panDB is especially powerful in classifying reads from high-complexity communities containing multiple conspecific strains and even unknown microbial species. Both databases can profile the majority of metagenomic data from human skin or stool microbiomes, with panDB also exhibiting significant sensitivity in characterizing intraspecific diversity. These database compilation pipelines can improve database-guided analyses of complex microbial communities by efficiently leveraging the rapidly expanding genome sequence data available in public databases.

## Methods

### Compilation of the representative genome database (reprDB)

To construct reprDB, we gathered genomes that maximize the representation of phylogenetically diverse microbes to reduce within-species redundancy. Representative and reference genomes, as curated and designated by PATRIC [29], were included in the database for each archaeal and bacterial species. Due to the heterogeneity of viruses, all viral genomes from NCBI’s accession list were included in the database [30]. Finally, fungal species were included from NCBI’s eukaryote genome browser.

UNIX shell scripts were developed to streamline database compilation. For each target organism, the FASTA file containing the genome sequence and a GenBank flat file containing the taxonomy of the organism were fetched using NCBI’s UNIX-compatible download tool E-utilities [31]. The FASTA file was then reformatted to encode the taxonomy and the genome size information in the header of the genome sequences. Genome sequences and their informative headers were concatenated into larger files of about 2.8GB, a convenient size for indexing by aligners such as Bowtie 2 [13]. The pipeline is especially suitable for parallel processing on a Portable Batch System. The pipeline is exceptionally user-friendly; the only required user inputs are 1) the name of the file that contains the target organisms, 2) the file format type (chosen from a provided list of compatible formats), and 3) the desired number of jobs generated by the script.

### Iterative whole-genome alignment

Briefly, in each iteration, a query genome is chosen and aligned to a standing reference genome (Figure 1B). The genome regions, or blocks, that are only present in the query genome but absent in the reference genome are identified and appended to the reference genome. In the next iteration, the updated reference genome is aligned with a different query genome to identify query-exclusive blocks, which are appended to the updated reference genome. The iteration continues until all strain genome sequences have been considered, either as a query genome or as the starting reference genome in the first iteration. Finally, the progressively updated reference genome sequence represents the species pan-genome sequence, which covers genomic regions in all strains. In the present study, the representative genome of a species was used as the reference genome in the first iteration. If a species does not have a designated representative genome, the genome with the greatest size was selected as the reference genome in the first iteration. The rest of the conspecific strain genomes were aligned iteratively to the reference genome in descending size order. We used Mugsy with default parameters for the alignment of conspecific strain genomes because Mugsy is especially suitable for closely related genome sequences [32].

### Compilation of the pan-genome database (panDB)

Automated panDB compilation was implemented in UNIX bash and C++, embedded in a UNIX shell wrapper, suitable for parallel processing on a Portable Batch System. All bacterial, fungal, archaeal, and viral genome assemblies were downloaded from GenBank and grouped by species, according to the species taxonomy ID provided by the assembly summary files [33]. The majority of microbial species only have one sequenced genome (Figure 8A, left panel), while the majority of sequenced genomes belong to species with multiple sequenced strains (Figure 8A, right panel). Within each species, genome sequences of different strains were aligned iteratively to identify the species pan-genome sequence. Some medically important species have hundreds or even thousands of sequenced genomes. In this study, to shorten the compilation time, only the representative genome and the 49 largest strain genomes were aligned for bacterial, archaeal and viral species that have more than 50 sequenced strain genomes. For fungal species that have more than 20 sequenced strain genomes, only the representative genome and the 19 largest strain genomes were used for the alignment. The maximum number of strain genomes used to compile the pan-genome sequence can be adjusted by the users. In this study, all contigs shorter than 1000bp, which constitute over 10% of the contigs (Figure 8B, left panel) but less than 1% of the bases in the database (Figure 8B, right panel), were removed from the final pan-genome sequences to further condense the database. Short contigs do not significantly improve read mapping due to poor contiguity, but can inflate space usage especially when long fasta headers are used. Users can specify the minimum contig length to keep when compiling panDB.

**Figure 8.**
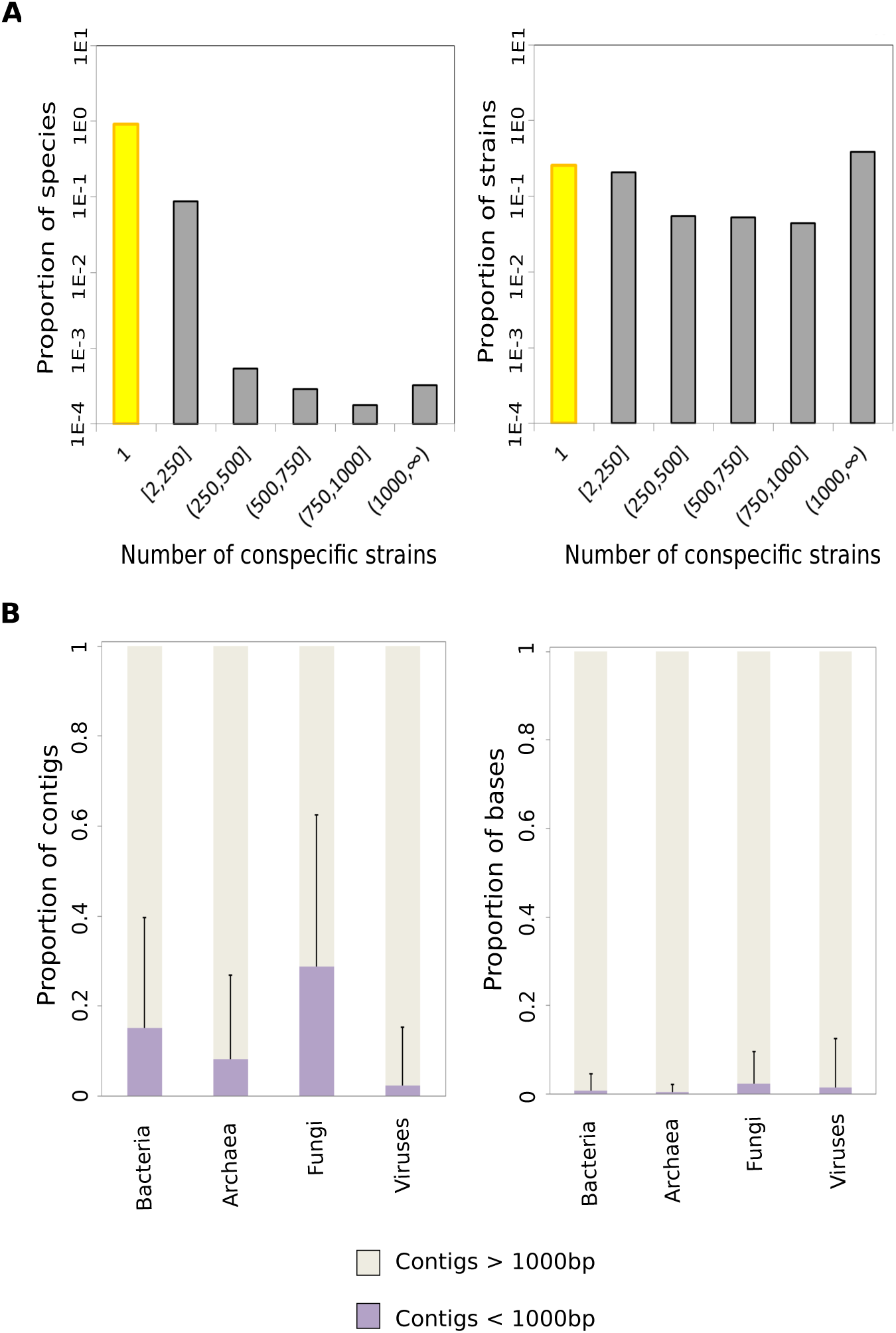
Length distribution of strain genomes and pan-genome contigs. 8A, distribution of the number of sequenced strains across microbial species in GenBank. Left panel, the proportion of species that have different numbers of sequenced strains. Right panel, the proportion of strains that have different numbers of sequenced conspecific strains. 8B, the length distribution of pan-genome contigs in panDB. Left panel, the proportion of short (less than 1000 bp) and long (greater than 1000 bp) contigs in the microbial pan-genome sequences. Right panel, the proportion of bases that are found within the short or long contigs.

### Detecting exogenous genomic sequences

We included a pipeline in the GitHub repository that detects potentially exogenous contigs from the pan-genome sequences in panDB. The pipeline extracts species- or strain-specific marker genes from the MetaPhlAn database [11], and searches the marker gene sequences against all pan-genome sequences in panDB using usearch-local [34]. The pipeline reports contigs that match any marker genes with sequence identities and e-values passing user-specified thresholds. The pipeline also reports the subset of contigs that either align to at least two marker genes, or align to at least one marker gene while the aligned region covering more than x% of the marker gene sequence, with the value of x specified by the users. In the present study, we used phyloT to visualize the species cladogram (http://phylot.biobyte.de/).

### Synthetic communities

To evaluate the sensitivity and specificity of the databases for read classification, we first used reprDB and panDB to analyze reads simulated from low-complexity *in silico* synthetic communities. To assess the sensitivity of the databases in recognizing multiple strains from the same species, synthetic communities were created with 5, 10, or 20 strain genomes of either *S. epidermidis* or *B. fragilis*. Next, the databases were tested for their sensitivity and specificity of species-level and phylum-level classification using synthetic communities created using representative genomes of 5 Bacteroides species (*B. fragilis, B. uniformis, B. vulgatus, B. ovatus, B. dorei*), 5 Staphylococcus species (*S. aureus, S. epidermidis, S. warneri, S. lugdunensis, S. capitis*), or 5 species representing 5 major bacterial phyla (*B. fragilis, S. epidermidis, Pseudomonas aeruginosa, Micrococcus luteus and Borrelia burgdorferi*). Assembly accession numbers of the genomes are available in Additional file 1. 500,000 or 1,000,000 Illumina reads were sampled from each of the 9 synthetic communities using Mason [35] with arguments –sq (simulating qualities), -i (include read information), -hs 0 (do not simulate haplotype snps), -hi 0 (do not simulate haplotype indels) and –n 100 (100bp read length). Read classification was conducted using Bowtie 2 [13] and Pathoscope 2.0 [9] as described below.

Next, to test the ability of the databases to classify reads from common bacterial species, we analyzed the mock metagenome community downloaded from ‘mockrobiota’ (mock community 17) [14, 15], which is the only shotgun metagenome dataset in mockrobiota and contains 21 evenly mixed bacterial strains. Low-quality bases were first trimmed using sickle with default parameters. The trimmed reads were then classified using Bowtie 2 [13] and Pathoscope 2.0 [9] as described below. The Pielou’s evenness index [36] was computed using:

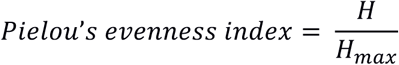

where *H* is the Shannon’s diversity index, and *H*_*max*_ is the maximum possible value of *H*:

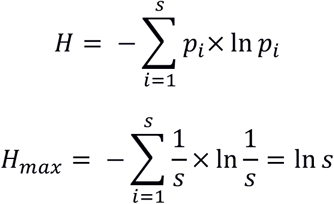

Where *p*_*i*_ is the relative abundance of the *i*th species, out of a total of *s* species. Finally, to test the ability of the databases to classify reads from unknown and high-complexity communities, we analyzed five high-complexity synthetic communities downloaded from CAMI (CAMI_high) [16], that were previously used in the first CAMI challenge. The datasets consist of 5 Hiseq samples of 15 Gbp each with small insert sizes sampled from complex synthetic communities containing over 700 predominantly unpublished isolate genomes. Read classification was conducted as described below, but due to the large size of the datasets, the reads aligned using Bowtie 2 [13] were not re-assigned using Pathoscope 2.0 [9]. Instead, the number of reads mapped to each genome were directly counted using samtools 1.5 [37].

### Construction of the Kraken databases and Kraken read classification

ReprDB and panDB were first formatted to include the NCBI taxID in their fasta headers according to the requirement by Kraken [7]. The databases were then built with hash-sizes of 10000M. We limited the maximum database sizes to be 256GB in order to fit the available memory space on our computer cluster. When a read can be mapped to multiple genomes, Kraken assigns the read to the lowest common ancestor of the genomes [7]. As a result, the more similar genomes a database contains, the more likely a read will map to multiple genomes and be assigned to a higher taxonomic level. Therefore, Kraken classification based on different databases cannot be compared on a single taxonomic level. Consequently, we compared the consistency between the taxonomic node a read is assigned to and the species genome from which the read is sampled – if the read is either assigned to the correct species or assigned to an ancestral node of the species, we conclude that the classification is consistent. We compared read assignment using three *in silico* synthetic communities generated in this study (communities with 5 *Bacteroides* species, 5 *Staphylococcus* species and 5 species representing 5 bacterial phyla, as described above) for which the species of origin of each simulated read is known. In addition, we assessed the sensitivity of read classification when multiple strains of a same species are present in a community. We did this by comparing the proportion of classifiable reads sampled from communities consisting of 5, 10 or 20 *S. epidermidis* or *B. fragilis* strains based on reprDB, panDB and the standard Kraken library.

### mWGS datasets

Using reprDB and panDB, we classified reads from 692 skin and 144 stool mWGS samples. The skin samples, as described previously in [4], were collected from 12 individuals from 17 defined anatomical skin sites (broadly classified as “moist”, “dry”, “sebaceous” and “foot”) over three time points. The stool samples, collected from 103 healthy adults, were acquired from the Human Microbiome Project (HMP) [17, 38].

### Read classification

The skin and stool mWGS datasets were first purged of low-quality reads and reads that mapped to the hg19 human reference as described [4]. The filtered reads were aligned to reprDB and panDB using Bowtie 2 version 2.2.9 under very-sensitive mode [13]. We rendered the aligner to look for at most 10 matches for each read in order to reduce computational time. Reads that align to more than one genome location were assigned to the most likely alignment using the PathoID module of Pathoscope 2.0 [9]. For taxonomic grouping, microbial species were grouped by their lowest taxonomic level at or above the genus level. Tentative classifications and misplaced classifications that are waiting for revisions were shown in square brackets according to the convention of the NCBI Taxonomy database [39]. Uncultured species—species that have no axenic culture for formal description [40]—were classified as an independent group. Taxonomic profiling of the stool samples based on 16s rRNA sequencing data using RDP classifier were downloaded directly from HMP (https://www.hmpdacc.org/hmp/HM16STR).

## Declarations

### Ethics approval and consent to participate

Not applicable

### Consent for publication

Not applicable

### Availability of data and materials

Database availability: The precompiled reprDB and panDB are available at: ftp://ftp.jax.org/zhouw/referenceDB/

### Software availability

Project name: reprDB, panDB Project home page: https://github.com/ohlab/reprDB, https://github.com/ohlab/panDB Operating system: Linux Programming language: UNIX shell, C++ Other requirements: Portable Batch System, GCC 4.9.2, Mugsy 1.2.3 License: GNU GPL v3.0

## Data availability

All reference genomes and simulated sequence reads from the *in silico* synthetic communities supporting the conclusions of this article are available upon request. The mockrobiota metagenome dataset is available from mockrobiota (https://github.com/caporaso-lab/mockrobiota/tree/master/data/mock-17) The 5 CAMI high-complexity datasets, the gold standard profiling, as well as the database used to generate the gold standard profiling are available from the first CAMI challenge (https://data.cami-challenge.org/participate) All human stool shotgun metagenomic data and 16s rRNA sequencing data supporting the conclusions of this article are available from HMP (http://hmpdacc.org/HMASM/) [17, 38] All human skin shotgun metagenomic data supporting the conclusions of this article are available in the NCBI Sequence Read Archive (bioproject 46333) [4, 41]

### Funding

This work was supported by National Institute of Health (K22 AI119231-01 to JO).

### Authors’ contributions

JO and WZ conceived the project. WZ and NG developed the algorithms and performed the analyses. JO and WZ drafted the manuscript. All authors read and approved the final manuscript.

### Acknowledgements

Not applicable

### Additional files

File name: Additional file 1.txt

File format: plain text (.txt)

Title of data: Assembly accession numbers of genomes used to generate the *in silico* synthetic community Description of data: a list of assembly accession numbers of genomes used to generate the *in silico* synthetic community

## References

1. Sharpton TJ. An introduction to the analysis of shotgun metagenomic data. Front Plant Sci. 2014; 5:209.

2. Qin J, Li R, Raes J, Arumugam M, Burgdorf KS, Manichanh C, Nielsen T, Pons N, Levenez F, Yamada T, et al. A human gut microbial gene catalogue established by metagenomic sequencing. Nature. 2010; 464:59–65.

3. Schloissnig S, Arumugam M, Sunagawa S, Mitreva M, Tap J, Zhu A, Waller A, Mende DR, Kultima JR, Martin J, et al. Genomic variation landscape of the human gut microbiome. Nature. 2013; 493:45–50.

4. Oh J, Byrd AL, Park M, Program NCS, Kong HH, Segre JA. Temporal Stability of the Human Skin Microbiome. Cell. 2016; 165:854–66.

5. Medini D, Donati C, Tettelin H, Masignani V, Rappuoli R. The microbial pan-genome. Curr Opin Genet Dev. 2005; 15:589–94.

6. Sunagawa S, Mende DR, Zeller G, Izquierdo-Carrasco F, Berger SA, Kultima JR, Coelho LP, Arumugam M, Tap J, Nielsen HB, et al. Metagenomic species profiling using universal phylogenetic marker genes. Nat Methods. 2013; 10:1196–9.

7. Wood DE, Salzberg SL. Kraken: ultrafast metagenomic sequence classification using exact alignments. Genome Biol. 2014; 15:R46.

8. Ames SK, Hysom DA, Gardner SN, Lloyd GS, Gokhale MB, Allen JE. Scalable metagenomic taxonomy classification using a reference genome database. Bioinformatics. 2013; 29:2253–60.

9. Hong C, Manimaran S, Shen Y, Perez-Rogers JF, Byrd AL, Castro-Nallar E, Crandall KA, Johnson WE. PathoScope 2.0: a complete computational framework for strain identification in environmental or clinical sequencing samples. Microbiome. 2014; 2:33.

10. Francis OE, Bendall M, Manimaran S, Hong C, Clement NL, Castro-Nallar E, Snell Q, Schaalje GB, Clement MJ, Crandall KA, Johnson WE. Pathoscope: species identification and strain attribution with unassembled sequencing data. Genome Res. 2013; 23:1721–9.

11. Segata N, Waldron L, Ballarini A, Narasimhan V, Jousson O, Huttenhower C. Metagenomic microbial community profiling using unique clade-specific marker genes. Nat Methods. 2012; 9:811–4.

12. Segata N, Boernigen D, Tickle TL, Morgan XC, Garrett WS, Huttenhower C. Computational meta'omics for microbial community studies. Mol Syst Biol. 2013; 9:666.

13. Langmead B, Salzberg SL. Fast gapped-read alignment with Bowtie 2. Nat Methods. 2012; 9:357–9.

14. Bokulich NA, Rideout JR, Mercurio WG, Shiffer A, Wolfe B, Maurice CF, Dutton RJ, Turnbaugh PJ, Knight R, Caporaso JG. mockrobiota: a Public Resource for Microbiome Bioinformatics Benchmarking. mSystems. 2016; 1.

15. Kozich JJ, Westcott SL, Baxter NT, Highlander SK, Schloss PD. Development of a dual-index sequencing strategy and curation pipeline for analyzing amplicon sequence data on the MiSeq Illumina sequencing platform. Appl Environ Microbiol. 2013; 79:5112–20.

16. Sczyrba A, Hofmann P, Belmann P, Koslicki D, Janssen S, Droege J, Gregor I, et al. Critical Assessment of Metagenome Interpretation - a benchmark of computational metagenomics software. bioRxiv. 2017; doi : 101101/099127.

17. Human Microbiome Project C. Structure, function and diversity of the healthy human microbiome. Nature. 2012; 486:207–14.

18. Wu D, Hugenholtz P, Mavromatis K, Pukall R, Dalin E, Ivanova NN, Kunin V, Goodwin L, Wu M, Tindall BJ, et al. A phylogeny-driven genomic encyclopaedia of Bacteria and Archaea. Nature. 2009; 462:1056–60.

19. Rinke C, Schwientek P, Sczyrba A, Ivanova NN, Anderson IJ, Cheng JF, Darling A, Malfatti S, Swan BK, Gies EA, et al. Insights into the phylogeny and coding potential of microbial dark matter. Nature. 2013; 499:431–7.

20. Shih PM, Wu D, Latifi A, Axen SD, Fewer DP, Talla E, Calteau A, Cai F, Tandeau de Marsac N, Rippka R, et al. Improving the coverage of the cyanobacterial phylum using diversity-driven genome sequencing. Proc Natl Acad Sci USA. 2013; 110:1053–8.

21. Lagier JC, Khelaifia S, Alou MT, Ndongo S, Dione N, Hugon P, Caputo A, Cadoret F, Traore SI, Seck EH, et al. Culture of previously uncultured members of the human gut microbiota by culturomics. Nat Microbiol. 2016; 1:16203.

22. Browne HP, Forster SC, Anonye BO, Kumar N, Neville BA, Stares MD, Goulding D, Lawley TD. Culturing of 'unculturable' human microbiota reveals novel taxa and extensive sporulation. Nature. 2016; 533:543–6.

23. Clingenpeel S, Clum A, Schwientek P, Rinke C, Woyke T. Reconstructing each cell's genome within complex microbial communities-dream or reality? Front Microbiol. 2014; 5:771.

24. Conlan S, Mijares LA, Program NCS, Becker J, Blakesley RW, Bouffard GG, Brooks S, Coleman H, Gupta J, Gurson N, et al. Staphylococcus epidermidis pan-genome sequence analysis reveals diversity of skin commensal and hospital infection-associated isolates. Genome Biol. 2012; 13:R64.

25. Jacobsen A, Hendriksen RS, Aaresturp FM, Ussery DW, Friis C. The Salmonella enterica pan-genome. Microb Ecol. 2011; 62:487–504.

26. Alneberg J, Bjarnason BS, de Bruijn I, Schirmer M, Quick J, Ijaz UZ, Lahti L, Loman NJ, Andersson AF, Quince C. Binning metagenomic contigs by coverage and composition. Nat Methods. 2014; 11:1144–6.

27. Scholz M, Ward DV, Pasolli E, Tolio T, Zolfo M, Asnicar F, Truong DT, Tett A, Morrow AL, Segata N. Strain-level microbial epidemiology and population genomics from shotgun metagenomics. Nat Methods. 2016; 13:435–8.

28. Nayfach S, Rodriguez-Mueller B, Garud N, Pollard KS. An integrated metagenomics pipeline for strain profiling reveals novel patterns of bacterial transmission and biogeography. Genome Res. 2016; 26:1612–25.

29. Snyder EE, Kampanya N, Lu J, Nordberg EK, Karur HR, Shukla M, Soneja J, Tian Y, Xue T, Yoo H, et al. PATRIC: the VBI PathoSystems Resource Integration Center. Nucleic Acids Res. 2007; 35:D401–6.

30. Brister JR, Ako-Adjei D, Bao Y, Blinkova O. NCBI viral genomes resource. Nucleic Acids Res. 2015; 43:D571–7.

31. Sayers E: E-utilities Quick Start. In Entrez Programming Utilities Help. Bethesda (MD); 2008

32. Angiuoli SV, Salzberg SL. Mugsy: fast multiple alignment of closely related whole genomes. Bioinformatics. 2011; 27:334–42.

33. Kitts PA, Church DM, Thibaud-Nissen F, Choi J, Hem V, Sapojnikov V, Smith RG, Tatusova T, Xiang C, Zherikov A, et al. Assembly: a resource for assembled genomes at NCBI. Nucleic Acids Res. 2016; 44:D73–80.

34. Edgar RC. Search and clustering orders of magnitude faster than BLAST. Bioinformatics. 2010; 26:2460–1.

35. Holtgrewe M. Mason – A Read Simulator for Second Generation Sequencing Data. FU Berlin. 2010.

36. Pielou EC. The measurement of diversity in different types of biological collections. J Theor Biol. 1966; 13:14.

37. Li H, Handsaker B, Wysoker A, Fennell T, Ruan J, Homer N, Marth G, Abecasis G, Durbin R, Genome Project Data Processing S. The Sequence Alignment/Map format and SAMtools. Bioinformatics. 2009; 25:2078–9.

38. Human Microbiome Project C. A framework for human microbiome research. Nature. 2012; 486:215–21

39. Federhen S. The NCBI Taxonomy database. Nucleic Acids Res. 2012; 40:D136–43.

40. Federhen S. Type material in the NCBI Taxonomy Database. Nucleic Acids Res. 2015; 43:D1086–98.

41. Oh J, Byrd AL, Deming C, Conlan S, Program NCS, Kong HH, Segre JA. Biogeography and individuality shape function in the human skin metagenome. Nature. 2014; 514:59–64.

